# Cross-scale Imaging of Live Botanical Specimens using Fourier Ptychographic Microscopy

**DOI:** 10.64898/2026.09.16.752051

**Authors:** Laura Copeland, Lewis D. Walker, Katherine Baxter, Mike Shaw, Gail McConnell

## Abstract

Light microscopic imaging of botanical specimens is typically performed using diffraction-limited methods which can image either a small field of view (FOV) with a shallow depth of field (DOF) at high resolution, or a large FOV and large DOF at low resolution. As such, many of the important small features, such as cells in the root cap of *Arabidopsis*, chloroplasts in algae and leaf guard cells, cannot be resolved in high numbers for meaningful statistical interpretation of results. Fourier ptychographic microscopy (FPM) is an accessible low-cost solution for large format bioimaging which makes use of multiangle illumination and iterative phase retrieval to recover high-resolution estimates of sample phase and amplitude. To date much FPM research has focused on optimising hardware and image reconstruction techniques with biological and biomedical applications limited primarily to mammalian cell cultures and histological tissue sections. We have, for the first time, applied FPM for label-free imaging of these large, living botanical specimens to investigate cellular and sub-cellular structures over multiple length scales simultaneously, with the aim of resolving structures that cannot be seen with diffraction-limited methods over a large field of view. With our FPM system we demonstrate imaging of living samples mounted in water with a half-pitch resolution of 615 nm over a 3.6 mm diagonal FOV, producing a measured 9-fold improvement in resolution compared to conventional brightfield imaging. This allows for individual root cap cells of *Arabidopsis thaliana* to be resolved within the context of larger root structures, individual chloroplasts identified in large section of *Spirogyra varians,* and high-resolution colour imaging of *Tradescantia zebrina* stomata and chloroplasts across large leaf sections.

## 1. Introduction

Imaging botanical specimens presents a unique challenge because structures of interest span multiple spatial scales, from whole organs to subcellular features. Conventional diffraction-limited microscopy requires a trade-off between spatial resolution and field of view (FOV): increasing numerical aperture (NA) improves resolution but simultaneously reduces both the FOV and depth of field (DOF).

Consequently, fine cellular structures can only be examined over limited regions and without their wider anatomical context. Although image tiling and stitching can extend the effective FOV, these approaches require long acquisition times and are susceptible to stage drift, sample movement and stitching artefacts. An alternative strategy is multiscale imaging, where images acquired at different magnifications are spatially aligned using image registration algorithms. While image registration is well established, existing methods have largely been developed for medical imaging and have not been extensively evaluated for botanical specimens. [1] *Arabidopsis thaliana* has been long established as a model organism in plant biology. [2] The roots of *Arabidopsis* plants have been used in studies of root development, responses to abiotic stresses, and microbial interactions. [3], [4], [5]*Spirogyra varians* is a filamentous freshwater alga with a distinctive spiral organisation of chloroplasts. Widely distributed in freshwater habitats worldwide, it has been applied in wastewater treatment to transform contaminants in the water found in places such as abandoned mining sites. [6] In rare cases, calcium oxalate crystals can form in *Spirogyra*. There is little known about the formation of these crystals and their function. *Tradescantia zebrina* is a variegated perennial plant native to Mexico. [7] Its leaves are used in traditional medicine for kidney ailments, treatment of high blood pressure, coughs and tuberculosis. [8] *Tradescantia* is also a popular specimen in educational environments, where it is used viewing stomata responses. [9] Its striking purple colouring comes from anthocyanin, a purple pigment that aids in light attenuation and light stress mitigation. [10], [11]

Diffraction-limited light microscopy is the preferred method of imaging these large, living specimens but forces a trade-off between FOV and spatial resolution This creates challenges when measuring features such as stomata over large volumes for statistical interpretation. It also prevents imaging of root cap cells within the context of larger root structures.

Fourier ptychographic microscopy (FPM) is a computational microscopy technique that combines angularly illuminated low-resolution intensity images to reconstruct high-resolution complex (amplitude and phase) images using iterative phase retrieval algorithms. [14, 15] Using a low-magnification objective lens and a light emitting diode (LED) array to provide the angular illumination, FPM captures a sequence of wide-field low-resolution intensity images that are computationally reconstructed into a single, high-resolution complex image. FPM has been successfully applied to blood smears, histological sections and cultured mammalian cells [12], [13], [14] but, to our knowledge, has not previously been demonstrated for imaging of living botanical applications, which are substantially thicker and more optically complex than specimens previously imaged with FPM. [15] Previous studies have found that due to the thin object assumption FPM algorithms are based upon, a maximum thickness of 10 μm is advised, however this has only been reviewed for histological samples and not botanical. [16]

The maximum half-pitch resolution limit of an FPM microscope is defined by rewriting the Abbe diffraction limit formula to the following. [15]

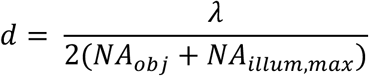

Where λ is the illumination wavelength, NA_obj_ is the NA of the objective lens and NA_illum,max_ is the maximum illumination NA, defined as *NA_illum_*_,*max*_ = *n sin*(*θ_max_*) with θ_max_ being the largest illumination angle. This implies that the shorter the illumination wavelength, the smaller the resolution of the image. FPM systems employ an RGB LED array for illumination, using approximately 630 nm for red, 530 nm for green and 470 nm for blue. It is expected that the resolution of the FPM images will vary with the different wavelengths of illumination, and that the greater thickness of the botanical specimens will accentuate differences between the reconstructed amplitude and phase images through increased phase wrapping and phase transfer characteristics.

Here, we investigate the application of FPM for imaging of living botanical specimens using three representative model systems, namely *Arabidopsis thaliana*, *Spirogyra varians, and Tradescantia zebrina*. We demonstrate high-resolution imaging across large fields of view, enabling visualisation of individual root cells, resolution of individual calcium oxalate crystals within algal filaments, and we perform multicolour imaging of pigmented leaf peels over a large FOV to establish the capabilities and limitations of FPM for plant imaging.

## 2. Methods

### 2.1. Botanical Specimen Growth and Preparation

*Arabidopsis thaliana* Col-0 (N1092, Nottingham *Arabidopsis* Stock Centre, UK) seeds were used. The seeds were sterilised in a bleach-based solution described in Supplementary Material and stored in the fridge for 2 days. They were then plated onto ½ Murashige and Skoog (MS) agar plates [17] and grown at 22°C in a 12hr light-dark cycle. The roots were then removed from the plant and mounted in 50 μL deionised water between a glass slide (TT-7041I-8217-001, Epredia, US) and a 24 × 50 mm type 1.5 coverslip (48393-172, VWR, US). The total sample size was n = 5 *Arabidopsis* plants, with 3 regions of interest (ROIs) imaged per plant, proving n = 15 image data sets.

*Spirogyra varians* (LZA075, Blades Biological Ltd, UK) was cultivated using 3N-BBM+V (Culture Collection of Algae & Protozoa, UK). The *Spirogyra* was mounted using 50 μL deionised water between a glass slide and 24 × 50 mm type 1.5 coverslip. The total sample size was n = 5 *Spirogyra* specimens with 3 ROIs imaged per specimen, proving n = 15 image data sets.

*Tradescantia* plants were grown in Peat-Free Multi-Purpose Compost (Ambassador, UK) indoors, with an average temperature of 20 °C and approximately 9 hours of indirect sunlight from an east-facing window. Mature leaves (approximately 4 weeks old) were selected and peeled using a scalpel to create a single cell layer and the peels were mounted using 20 μL deionised water between a glass slide and 22 × 22 mm type 1.5 coverslip (631-0124, VWR, USA). The total sample size was n = 10 leaf peels from 4 different leaves.

Coverslip sizes varied depending on the size of the sample, the *Arabidopsis* roots and *Spirogyra* filaments required the larger 24 × 50 mm coverslips whereas the *Tradescantia* leaf peels were smaller, requiring the 22 × 22 mm coverslips.

### 2.2. FPM Imaging and Data Acquisition

Imaging was performed using a home-built horizontal FPM set-up for ease of alignment of the LED array and optical components. A schematic diagram of this set-up is shown in Figure 1. The illumination source was a low-cost consumer-grade RGB LED array (WS2812B, BTF Lighting, UK) with wavelengths λ_Red_ = 636 nm, λ_Green_ = 523 nm, and λ_Blue_ = 470 nm and an LED pitch of 10mm. The array was selected due to its low cost and compatibility with a Raspberry Pi 4 model B, which used an open-source Python-based single-script graphical user interface to control the LED array and create/ use preloaded patterns for imaging. [18] A Nikon 4X 0.13 NA objective lens was used in combination with a 200 mm focal length tube lens (TTL200-A, Thorlabs, Germany) together with a monochromatic iDS UI-3060CP-M-

**Figure 1:**
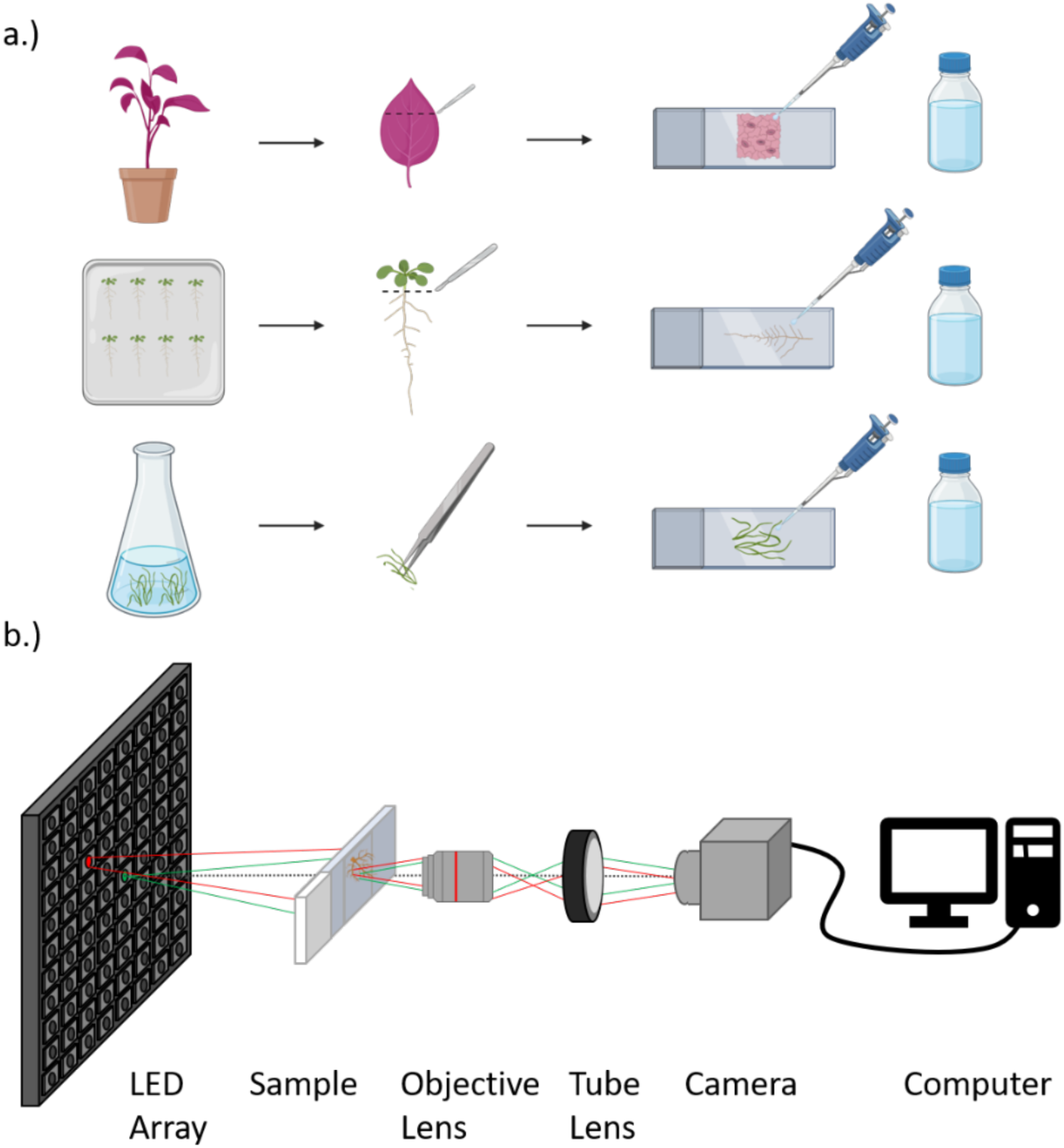
a.) Growth conditions and sample preparation for the 3 botanical specimens. *Tradescantia zebrina* was grown on compost, and a single cell layer was peeled with a scalpel before mounting between a slide and coverslip with water. *Arabidopsis thaliana* was grown on ½ MS agar, the roots were detached using a scalpel and specimens were mounted between a slide and coverslip with water. *Spirogyra* was grown in in 3N-BBM+V growth media, and filaments were collected using tweezers and mounted between a slide and coverslip with water. b.) FPM optical diagram with LED array pattern of LEDs. The green LED and light path indicates a brightfield image/LEDs and the red LED and light path represent a darkfield image/LED.

GL R2 camera for detection and image formation, using software (ThorCam 64-Bit Windows version 3.7.0) to collect data. [19] The samples were mounted using an XY translation mount (XYF1, Thorlabs, Germany), mounted on a single-axis translational stage with standard micrometre used for focusing the sample (PT1, Thorlabs, Germany).

For all specimens, 177 LEDs were activated sequentially in a circular pattern, and the corresponding 177 images were collected and saved as a TIFF stack. For the monochrome images, the red LEDs were used with an exposure time of 90 ms. The total capture time of 177 images was approximately 90 s. To create a brightfield image that was comparable to conventional brightfield microscopy, the 177 raw images were summed by performing a summed slices z projection in FIJI [20] to make an 8-bit incoherent image. The images all had the same bit depth (8-bit), but file sizes varied as the incoherent images were 9.2 MB whereas the FPM amplitude and FPM phase images were 258 MB. As a result, the FPM images were divided into 6 tiles for further processing as described in Section 2.4.

For the FPM reconstructions, a sequential quasi-Newton phase retrieval algorithm [21] was implemented in the open-source FPM app written by Rogalski et al. [22], [23]. The PC used for the reconstruction algorithm was equipped with Intel^®^ Xeon^®^ E-2124G CPU (3.4 GHz 64 GB RAM). During the reconstruction process raw images (1936 pixels × 1216 pixels × 177 frames) were divided into 28 tiles (2400 pixels × 2400 pixels) with a 10% overlap. This reduced the computational memory limitations [24] and satisfied the spatial coherence constraints imposed by the van Cittert-Zernike theorem. [25] Tiles were then stitched together using the Grid/Collection stitching plugin on FIJI to create the full FOV. [20]

For colour imaging, a sample dataset was collected in each of the 3 colour channels using the red, green, and blue LEDs with exposure times of 90 ms, 90 ms, and 50 ms respectively. For the colour incoherent image the summed red, green and blue incoherent images were merged using the colour merge channels tool on FIJI. [20] For the colour FPM amplitude images each of the red, green, and blue FPM amplitude images were reconstructed separately and then lateral chromatic offsets were corrected using rigid registration [26] and the 3D drift correct plugin on FIJI. [27] The corrected FPM amplitude red, green, and blue images were then merged using the colour merge channels tool on FIJI to create the colour FPM amplitude image.

In an FPM reconstructed phase image, phase values are assigned a value between - π and π, However, in practice, parameters such as the illumination wavelength, the refractive indices of the sample and surrounding medium, and the sample thickness contribute to the accumulated phase shift. Consequently, optically thick samples (> 10 *μm*) [16], [23] or regions with steep variations in thickness or refractive index can produce phase shifts that exceed 2π. [28] Because the reconstructed phase is represented as modulo 2π, phase values extending beyond this limit are mapped back into this range, producing phase discontinuities known as phase wrapping.

Furthermore, if the spatial phase change between adjacent pixels exceeds π radians, the phase becomes insufficiently sampled, making the number of 2π phase cycles ambiguous and prevents reliable unwrapping. A phase unwrapping algorithm is used to achieve this remapping by resolving the discontinuities into a continuous phase function. [29] Phase unwrapping of the *Spirogyra* samples was performed using the MATLAB algorithm written by Herráez et al. [30] The algorithm uses a non-continuous path to unwrap points with higher reliability values.

#### Synthetic Numerical Aperture and Resolution Measurements

The synthetic NA (NA_syn_) of an FPM system is described as *NA_syn_* = *NA_ob_*_j_ + *NA_illum_*_,*max*_ [28]. With the distance between the sample and LED array set to 121 mm, an LED pitch of 10 mm and 15 LED diameter circle, the NA_illum,max_ was calculated to be 0.52, giving a synthetic NA of 0.65. This value was then substituted 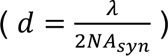) providing a theoretical spatial half-pitch resolution of 489 nm at *λ* = 636nm. A United States Air Force (USAF) test target (2015a, Ready Optics, Calabasas, CA, USA) was imaged to measure the experimental resolution limit of the system, and therefore the experimental synthetic NA. A line plot profile was plotted on FIJI and the smallest resolvable line pairs were compared to the USAF target data sheet from the supplier. [Supplementary material] [29]

### 2.3. Specimen Dimension Measurements

*Spirogyra* diameters were measured using the line tool in FIJI along three different sections of the length of the filament. These measurements were averaged and the error taken was the standard deviation.

### 2.4. Image Decorrelation Analysis

To investigate how the resolution varied with different botanical samples and illumination wavelengths, the resolution of the incoherent brightfield, FPM amplitude, and FPM phase images were measured using image decorrelation analysis for the *Arabidopsis*, *Spirogyra* and *Tradescantia* including each colour channel for the *Tradescantia* colour images. [33] Three datasets were analysed per specimen with 3 colour channels for each of the image types for the *Tradescantia* images. As the FPM images have large file sizes, the FOV was divided into 6 ROIs, and each ROI was analysed on FIJI using the image decorrelation analysis plugin using a maximum area of 0.5. The resolution for each image was then taken from these results as an average across the 6 ROIs. Supplementary materials show the resulting decorrelation curves, including the amplitude of the local maximum of the decorrelation function, *A_0_*, and the calculated cut-off frequency, *k_c_*. For the *Arabidopsis* and *Spirogyra* data the resolution of the incoherent brightfield is presented as an average of n = 3 datasets, and the resolution of the FPM amplitude and FPM phase images are presented as an average of n = 6 ROIs across n = 3 datasets. For the *Tradescantia* data resolution for the incoherent brightfield is presented as an average of n = 3 datasets for each wavelength of LED, whilst the resolution of the FPM amplitude and phase are presented as an average of n = 6 ROIs across n = 3 datasets.

## 3. Results

### 3.1. FPM imaging of *Arabidopsis* roots

Figure 2 shows representative images of overlapping mature *Arabidopsis* lateral roots imaged with the FPM system. Due to the large FOV, it was possible to image large root segments that included the root cap, apical meristem, elongation zone and maturation zone with cellular resolution. Figure 2 a.) shows the full FOV divided into 3 to show the incoherent brightfield image (left), the FPM amplitude image (centre), and the FPM phase image (right). The cyan and red boxes display digital zoom ROIs of the incoherent brightfield, FPM amplitude and FPM phase images, with figures 2 b.) – 2 d.) showing the lateral root cap, and figures 2 e.) - 2 f.) showing two overlapping lateral roots. Phase wrapping artefacts occur in 2 a.) (right), d.) and f.), results following the use of an unwrapping algorithm are shown in the supplementary material.

**Figure 2:**
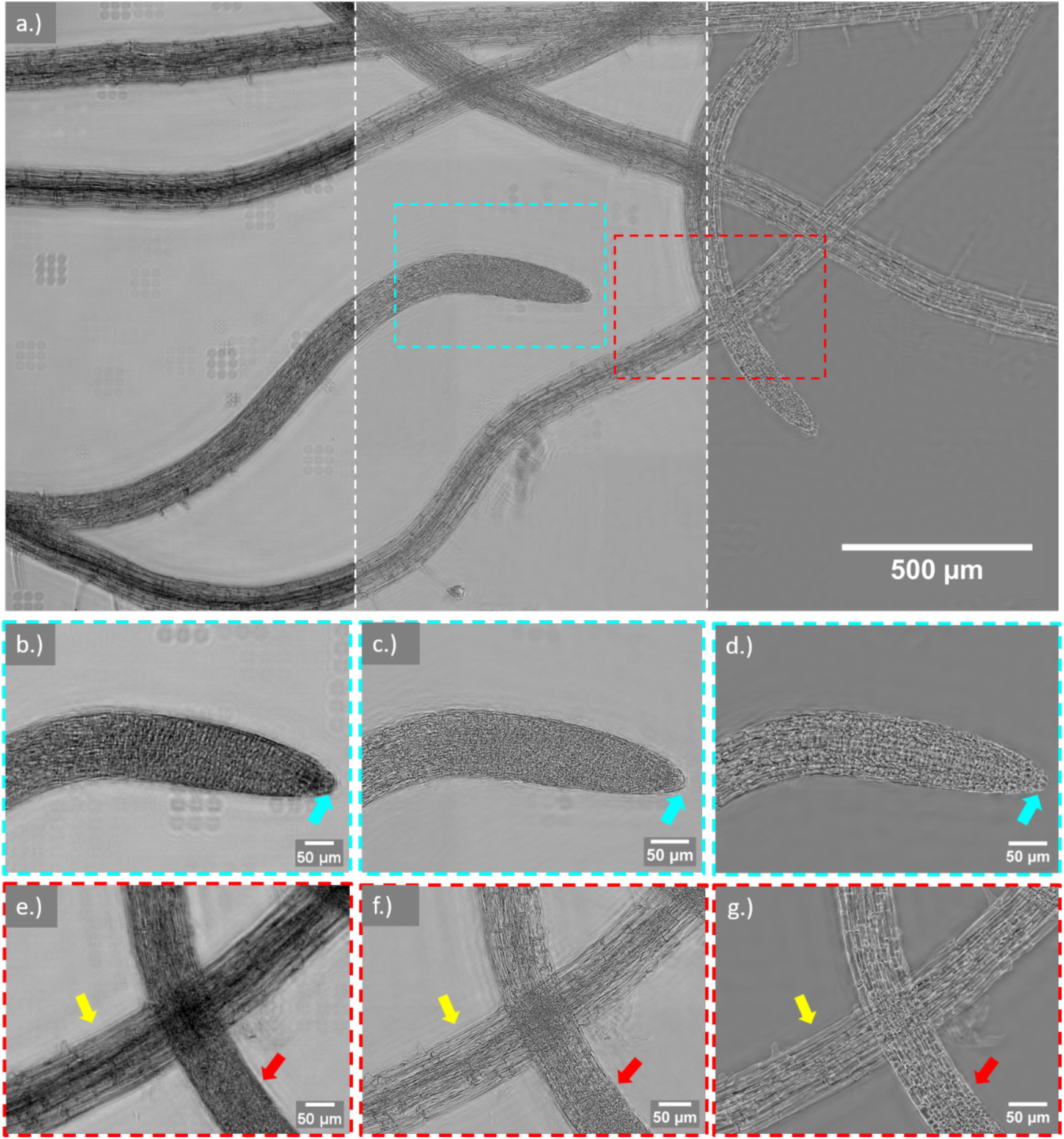
*Arabidopsis* root: a.) The full FOV has been divided into 3 to show the incoherent brightfield image (left), reconstructed FPM amplitude image (middle) and the reconstructed FPM phase image (right). Both the brightfield and reconstructed FPM amplitude images have been contrast adjusted using FIJI. [20] Cyan and red boxes highlight digital zoom ROIs. ROIs of the incoherent brightfield images are shown in b.) and e.), ROIs of the FPM intensity image are shown in c.) and f.), while the ROIs of the FPM phase image are shown in d.) and g.). The cyan boxes show ROIs of the root tip with the cyan arrow highlighting the root cap cells that are not seen in the incoherent brightfield image but are clearly seen in the FPM intensity and phase images. The red boxes show two overlapping lateral roots. The red arrow indicates that one lateral root is in a different focal plane to a second lateral root indicated with the yellow arrow. In the FPM reconstruction images cellular details are resolved in both lateral roots, despite the overlap.

As the incoherent brightfield image is a sum of all 177 images in an FPM dataset, out of focus features are shifted laterally by the varied illumination angle in each raw coherent brightfield image. This results in the 9 dust spots arranged in a grid that can be seen in the left part of figure 2 a.). An advantage of FPM is that the increased NA, and correspondingly reduced DOF, suppresses the appearance out of focus information in each image showing only the part of the specimen within that DOF.

Using FPM, the complex structures in two overlapping roots can be resolved. In figures 2 e.) – g.) 2 lateral roots cross, with the root (indicated by red arrows) appearing out of focus in the incoherent image (figure 2 e.)). Despite this, the reconstruction algorithm can detect structure and produce FPM amplitude and FPM phase images allowing individual cells and the internal vasculature to be resolved.

#### 3.1.1. Resolution measurement of *Arabidopsis*

Figure 3 shows the results of using image decorrelation analysis. The resolution of the incoherent brightfield, FPM amplitude and FPM phase images were measured to be 15.3 ± 9.21 μm, 0.82 ± 0.1 μm and 1.20 ± 0.37 μm respectively. A Mann-Whitney test showed statistically significant difference in resolution between image types, with p = 0.008, p < 0.0001, and p = 0.0015 for amplitude – incoherent, amplitude – phase and incoherent – phase, respectively.

**Figure 3:**
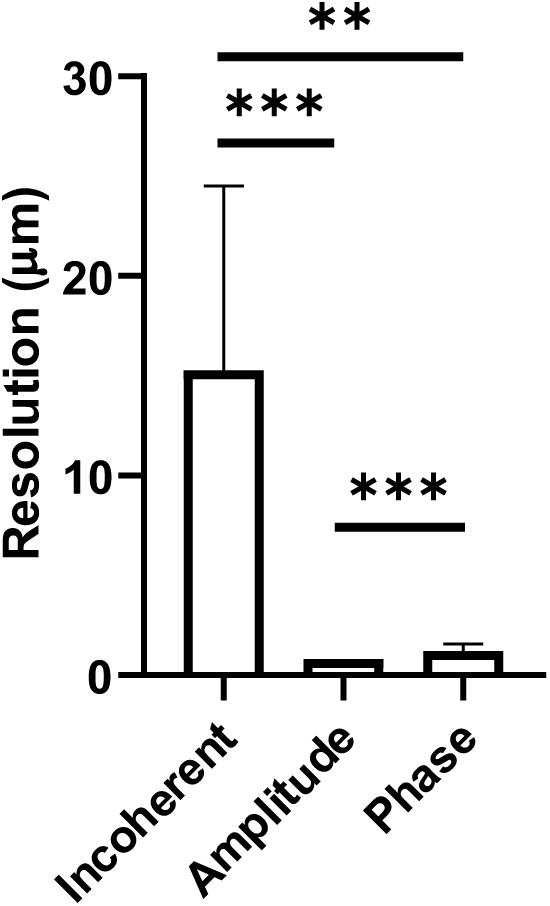
Results of image decorrelation analysis of *Arabidopsis* thaliana images.

### 3.2. FPM imaging of *Spirogyra* algae

Figure 4 shows representative FPM images of *Spirogyra varians*. Figure 4 a.) is the full FOV incoherent brightfield image, 4 b.) is the full FOV FPM amplitude image, 4 b.) and d.) show full FOV FPM phase images before and after phase unwrapping. The average diameter of the *Spirogyra* specimens was measured to be 180.06 ± 3.9 μm, which is thicker than most specimens studied with FPM to date. Digital zoom ROIs of the incoherent brightfield image, FPM amplitude image, wrapped FPM phase image, and unwrapped FPM phase image are displayed in figures 4 e.) to h.) respectively. The ROIs highlight structures such as chloroplasts and calcium oxalate crystals that cannot be seen in the incoherent brightfield images but are clearly visible in the reconstructed FPM images.

**Figure 4:**
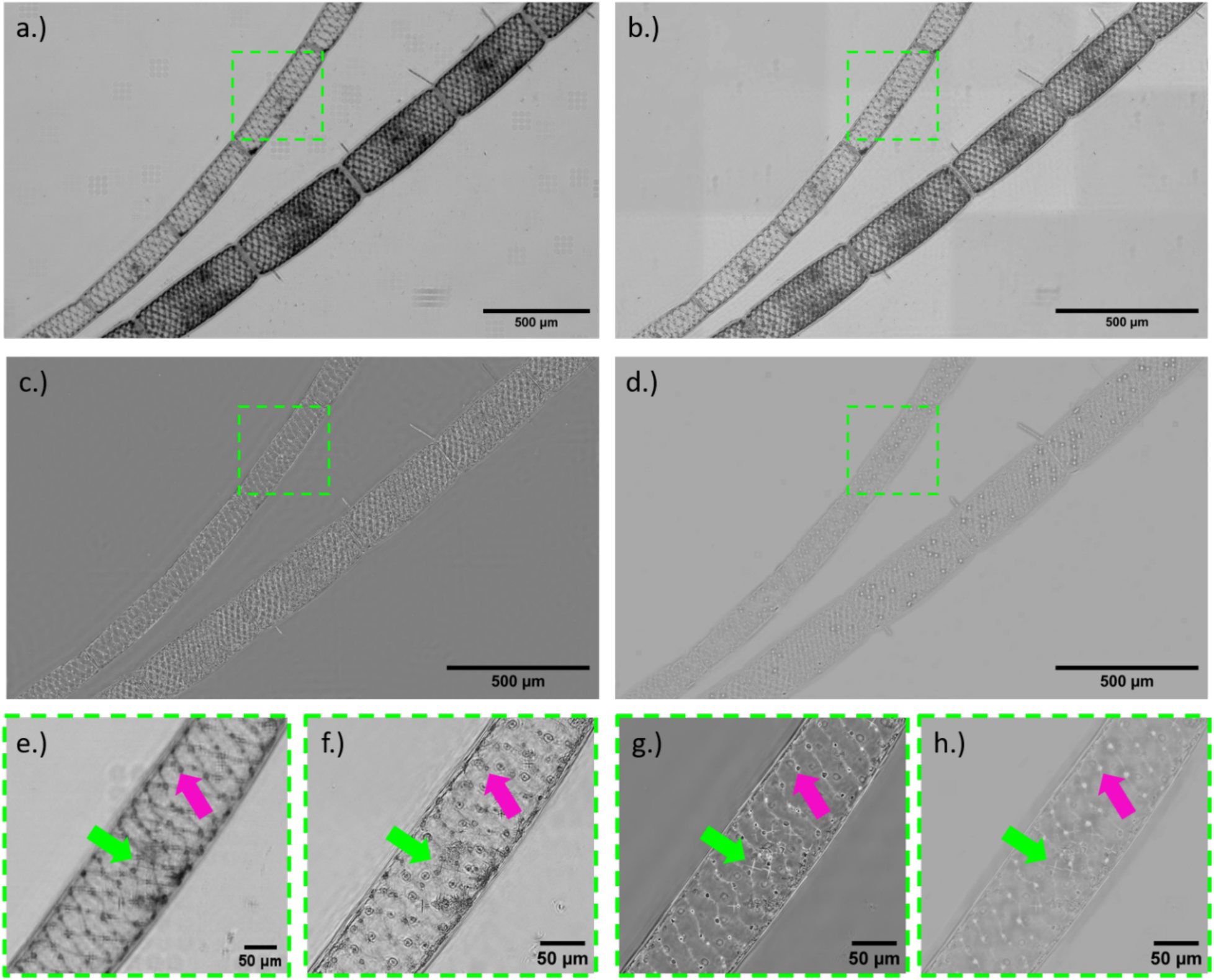
*Spirogyra varians*: a.) Incoherent brightfield, contrast adjusted on FIJI. [20] b.) FPM Amplitude image, contrast adjusted on FIJI. c.) FPM phase with wrapping artefacts. d.) FPM phase after un-wrapping, and with CLAHE. e.), f.), g.) and h.) are digital zoom ROIs of a.), b.), c.) and d.) respectively. The FPM outputs show clearly resolved individual chloroplasts, highlighted with the magenta arrow, arranged in the spiral that *Spirogyra* is named for and calcium oxalate crystals, highlighted with the green arrow.

#### 3.2.1. Resolution measurement of *Spirogyra*

Figure 5 shows the results of the image decorrelation analysis, the resolution of the incoherent brightfield, FPM amplitude and FPM phase images were measured to be 7.46 ± 1.03 μm, 0.81 ± 0.1 μm and 1.10 ± 0.39 μm respectively. A Mann-Whitney test showed statistically significant difference in resolution between image types, with p = 0.008, p = 0.0003, and p = 0.0015 for amplitude – incoherent, amplitude – phase and incoherent – phase, respectively.

**Figure 5:**
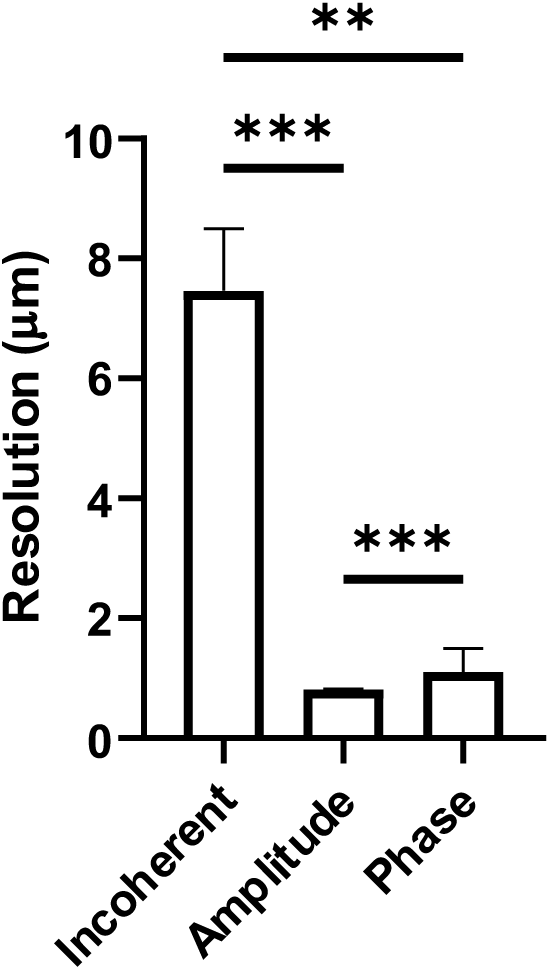
Results of the image decorrelation analysis of *Spirogyra varians* images.

### 3.3. FPM imaging of *Tradescantia zebrina*

Figure 6 a.) shows representative full FOV colour imaging with FPM. The top right is a merge of the incoherent brightfield images illuminated with red, green, and blue LEDs, whilst the bottom left is a merge of the FPM amplitude images illuminated with red, green, and blue LEDs. Digital zoom ROIs in figures 6 b.) and c.) show a stomata, guard cells and surrounding subsidiary cells in the incoherent brightfield and FPM amplitude images respectively. In the FPM amplitude it is possible to distinguish indivdual leucoplasts surrounding the nucleus of each cell across hundreds of cells in the full FOV that are not clearly visible in the brightfield image.

**Figure 6:**
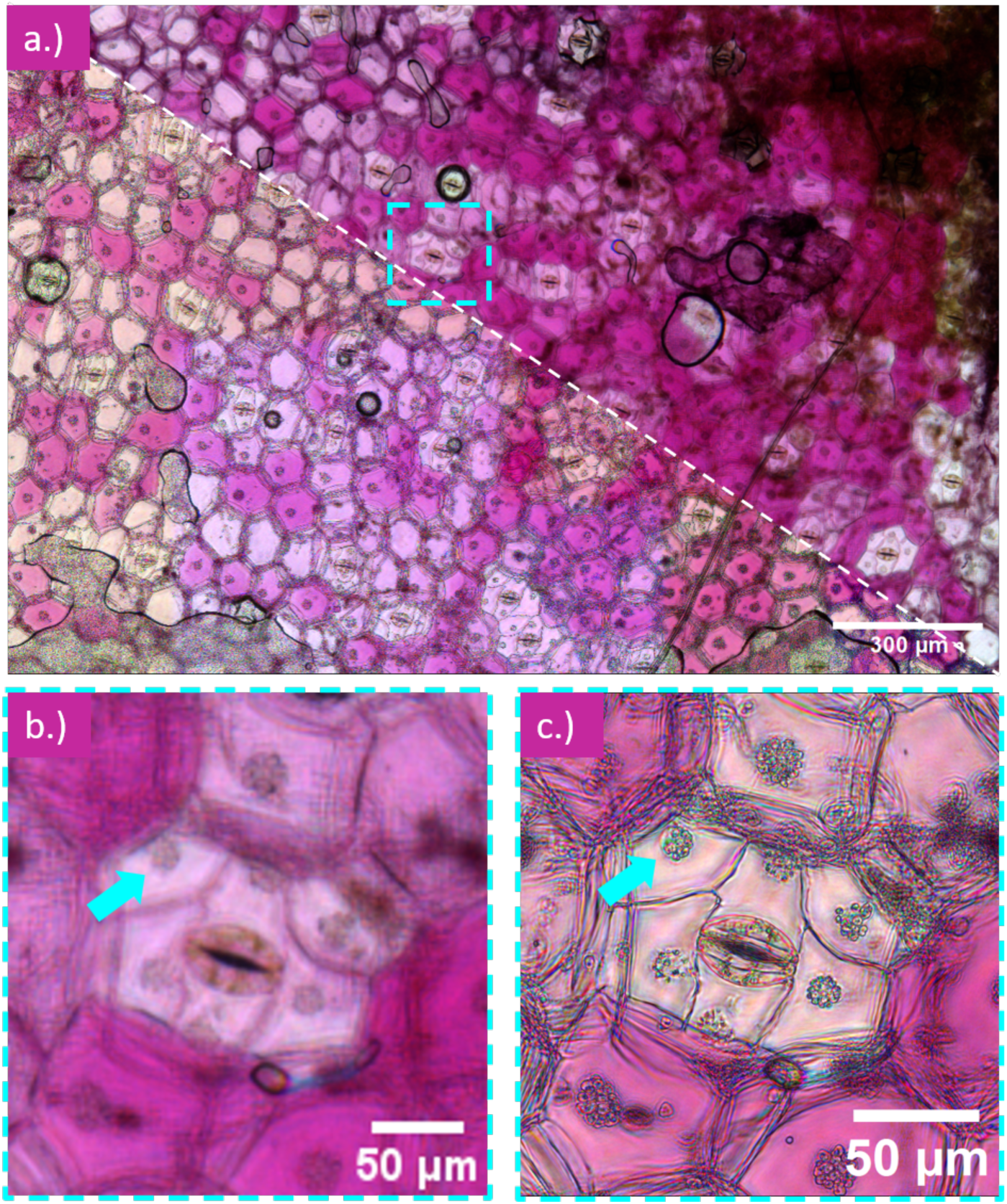
*Tradescantia zebrina*: a.) full FOV is divided diagonally to show the incoherent brightfield (top right) and FPM amplitude (bottom left). b.) and c.) are ROIs of the incoherent brightfield and the FPM amplitude images respectively, a cyan arrow highlights a single leucoplast surrounding the cell nucleus that can be seen in the FPM amplitude image but not clearly in the incoherent brightfield.

### 3.4. Colour Resolution Measurements

Using a USAF test target and red illumination (636 nm), the half-pitch resolution and synthetic NA were measured to be *d* = 615 nm (Group 9 Element 5) and *NA_syn_* = 0.52, respectively. [Supplementary material] Figure 7 shows the results of the image decorrelation analysis. Figure 7 a.) shows the resolution results of the incoherent brightfield images for each wavelength, a Mann-Whitney test showed no statistical significance between any of the wavelengths with a significance of p > 0.999 for all. Figure 7 b.) shows the resolution results for the FPM amplitude images for each wavelength, a Mann-Whitney test showed no statistical significance between any of the wavelengths. p = 0.67, p = 0.23 and p = 0.27 for red – green, red - blue and green – blue, respectively. Figure 7 c.) is the resolution results of the FPM phase images for each wavelength, a Mann-Whitney test showed no statistical significance between any of the wavelengths. p = 0.58, p = 0.76, and p = 0.53 for red – green, red – blue, and green – blue, respectively.

**Figure 7:**
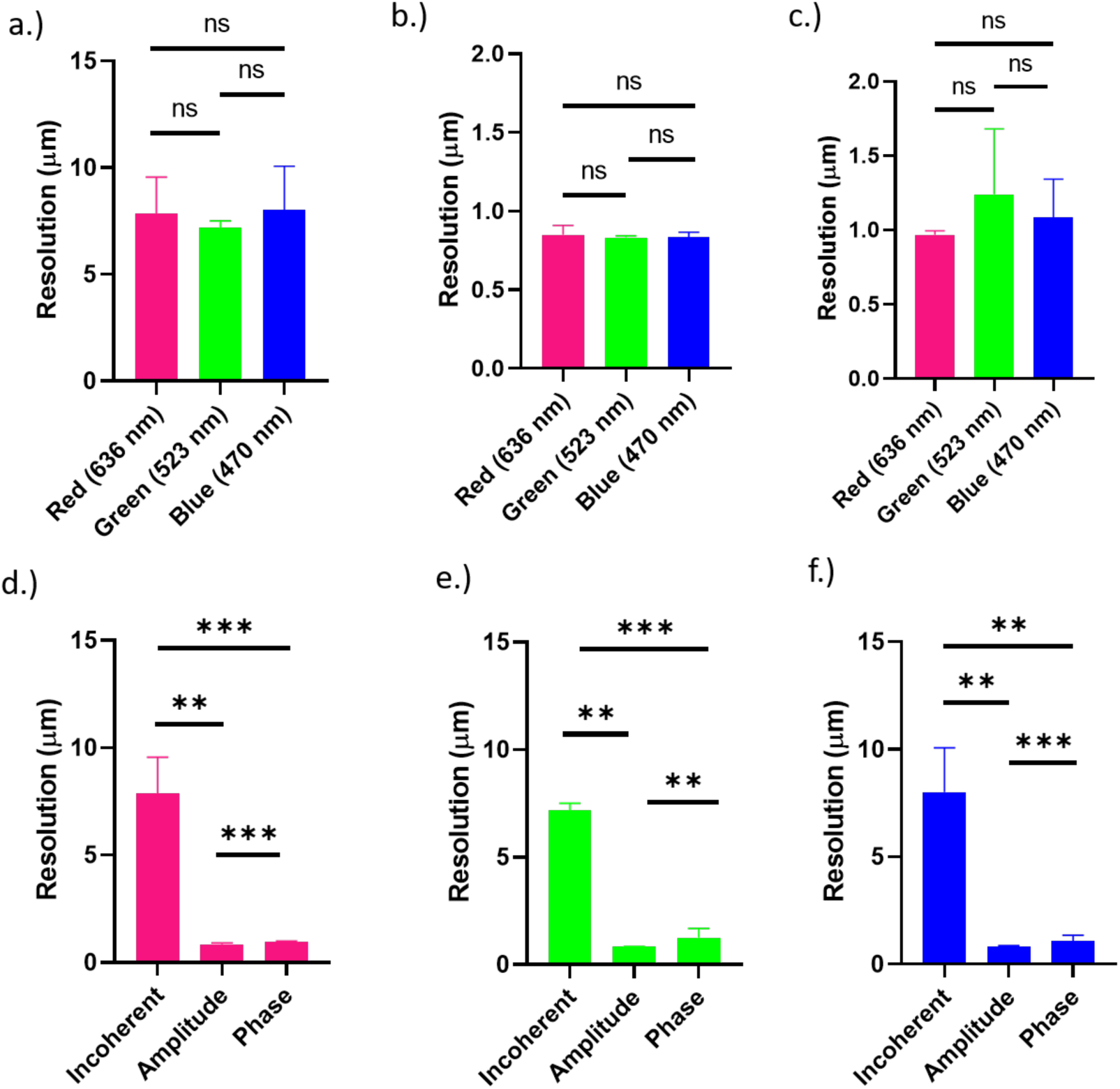
Resolution measurements from image decorrelation analysis: a.) resolution from the incoherent images at three different wavelengths red (636 nm), green (523 nm) and blue (470 nm) in magenta, green and blue respectively. b.) results from the FPM amplitude images and c.) is results from the FPM phase images. d.) results from imaging with 636 nm LEDs, e.) results from imaging with 523 nm LEDs and f.) results from imaging with 470 nm LEDs.

Figures 7 d.) – f.) show the resolution results grouped by each wavelength for each of the image types. Using Mann-Whitney tests between each variable, the results show statistical significance between resolution measurements depending on image type. For the images produced by the red LEDs (magenta plot), the p values were p = 0.002, p < 0.0001, p = 0.0009 for amplitude – incoherent, amplitude – phase and phase – incoherent respectively. For the images produced by the green LEDs (green plot), the p values were p = 0.0015, p = 0.002, p = 0.0008 for amplitude – incoherent, amplitude – phase and phase – incoherent respectively. For the images produced by the blue LEDs (blue plot), the p values were p = 0.001, p < 0.0001, p = 0.0015 for amplitude – incoherent, amplitude – phase and phase – incoherent respectively.

The resolution measurements show that FPM provides a 9-fold resolution improvement for amplitude images compared to the conventional brightfield imaging.

This result remains consistent across all three illumination wavelengths. This allows for visualisation of small features such as leucoplasts in guard cells over a large FOV.

## 4. Discussion

This study demonstrates that Fourier ptychographic microscopy can be successfully applied to living botanical specimens spanning a wide range of morphologies and thicknesses. Across *Arabidopsis thaliana* roots, *Spirogyra varians* filaments and *Tradescantia zebrina* leaf peels, FPM consistently provided higher spatial resolution than conventional incoherent brightfield imaging while maintaining a millimetre-scale field of view. Furthermore, multicolour FPM imaging was achieved without measurable differences in reconstructed resolution between illumination wavelengths.

When imaging *Arabidopsis thaliana* roots the increased spatial resolution was measured in both the FPM amplitude (0.82 ± 0.01 μm) and FPM phase (1.20 ± 0.37 μm) images as compared to the brightfield incoherent image (15.3 ± 9.21 μm). This allowed for visualisation of individual root cells and hairs over large root sections, including the elongation zone, meristematic zone and root cap within one FOV. Moreover, the reduced depth of field allowed for improved sectioning and removal of out of focus material, allowing for reconstructions of overlapping lateral roots.

By imaging *Spirogyra varians* with FPM it was possible to resolve subcellular details such as the organisation of chloroplasts and calcium oxalate crystals across a large FOV. *Spirogyra* was the thickest of the specimens imaged, with an average diameter of 180.06 ± 3.9 μm. Despite the thin object assumption, the FPM algorithm produced high resolution amplitude and phase images, with a 9-fold resolution improvement between the incoherent brightfield images and the FPM amplitude images. This suggests that FPM algorithms are more tolerant of thicker specimens and that the density of the specimen has an impact on the phase retrieval algorithm than sample thickness alone. This is demonstrated in attempts to remove phase wrapping artefacts in the *Arabidopsis* images. The cellular organisation in the *Arabidopsis* root tip is more dense than the *Spirogyra* filaments and attempts to use the phase unwrapping algorithm for the *Arabidopsis* roots resulted in further image artefacts. [Supplementary material] FPM amplitude reconstructions remained visually robust despite increased specimen thickness, whereas phase reconstructions degraded owing to multiple 2π phase discontinuities and increased optical path length variation. These observations suggest that amplitude imaging may currently represent the more reliable output of FPM for thick botanical specimens, while further advances in phase retrieval and unwrapping algorithms will be required before quantitative phase imaging using FPM becomes equally robust.

Other sample dependent artefacts were seen due to reconstruction algorithm working in patches. In samples such as Figure 3 b.) where there were entire patches that have no complex information from a biological sample, the reconstruction algorithm did not have a point of comparison for contrast and therefore, the background was generated darker than the background of tiles with sections of *Spirogyra*. Future work aims to remove these patches and create a more homogenous background for both the amplitude and phase images to facilitate more accurate object detection and segmentation for downstream analysis.

We have demonstrated that FPM systems are capable of high resolution colour imaging of botanical samples through RGB LED illumination, to image *Tradescantia* leaf peels. Image decorrelation analysis revealed no statistically significant difference in FPM image resolution between the 3 illumination wavelengths. This result was unexpected as based on the revised Abbe diffraction limit formula it was expected that the shorter wavelength would result in improved resolution. Nevertheless, there was statistically significant difference between the resolution of the incoherent brightfield, the FPM amplitude and the FPM phase images. The difference in resolution between the FPM amplitude and FPM phase images is likely due to phase wrapping artefacts in the FPM phase images.

This work shows the potential of FPM as a tool in botanical labs. Time-lapse FPM would allow for studies of *Arabidopsis* root hair growth over large FOVs, or how changes to environment alters calcium oxalate crystal formation in *Spirogyra*. The colour imaging of pigmented samples such as *Tradescantia* would allow for more detailed studies of anthocyanin localization over statistically large areas. In addition, there has been research into further reducing the cost of FPM systems and to create more portable microscopes. [31], [32], [33] This would provide a low-cost solution to high resolution, large FOV imaging of botanical samples in more remote research stations or low resource settings.

Overall, this work demonstrates that FPM extends beyond its traditional applications of thin mammalian cell and tissue samples and can successfully image living botanical specimens that are substantially thicker and structurally more complex. By combining sub-micron spatial resolution with millimetre-scale fields of view using low-cost hardware, FPM provides a promising platform for quantitative plant imaging in both laboratory and field environments.

## Author Contributions

**Laura Copeland**: Methodology; investigation; resources; writing – original draft; conceptualisation. **Lewis D. Walker**: Resources; writing – review and editing. **Katherine Baxter:** Resources; writing-review and editing. **Mike Shaw:** writing – review and editing; conceptualisation; supervision; funding acquisition. **Gail McConnell:** writing – review and editing; conceptualisation; supervision; funding acquisition.

## Acknowledgements

We wish to thank Dr Joel Milner’s group (University of Glasgow, UK) for use of laboratory space and equipment for the growth of *Arabidopsis thaliana*.

## Funding

LC and LDW were supported by the Engineering and Physical Sciences Research Council (EP/Y528833/1) and UK Government’s Department for Science, Innovation & Technology through the National Measurement System. KB was supported by the Biotechnology and Biological Sciences Research Council BB/Z516120/1]. MS acknowledges funding from the UK Government’s Department for Science, innovation & Technology through the Life Sciences and Health Program of the National Measurement System and the NIHR University College London Hospitals Biomedical Research Centre (award 187809). GM was supported by the Leverhulme Trust and supported in part by the Biotechnology and Biological Sciences Research Council (BB/V019643/1, BB/X005178/1, BB/T011602/1, and BB/W019032/1).

## Disclosures

The authors declare no conflict of interest.

## Data Availability Statement

The data presented in this article are publicly available on request from the University of Strathclyde Knowledge-Base at https://doi.org/10.15129/0d7cf18c-0024-4a31-9f40-18ec45593122. This dataset includes raw images in .tiff format for all imaging and the results from the image decorrelation analysis.

## Notes

https://doi.org/10.15129/0d7cf18c-0024-4a31-9f40-18ec45593122

### Competing Interest Statement

The authors have declared no competing interest.

### Summary of Updates

Small typo from mm to μm for one of the resolution results.

