## Supplementary Material for "Cross-scale Imaging of Live Botanical Specimens using Fourier Ptychographic Microscopy"

**Seed Sterilisation:**

- 35ml dH_2_O
- 4 drops triton X-100
- 1 chlorine tablet
- 45 ml 100% EtOH
- 50 ml 70 % EtOH
- 50 ml sterile dH_2_O

Sterilising solution:

Add the 4 drops of triton X-100 to the 35 ml dH_2_O, place in the fume hood and add the chlorine tablet. Leave until dissolved.

Add 5 ml of above solution into 45 ml of 100% EtOH and centrifuge at 3500 rpm for 5 mins at 20 °C.

Place seeds in an Eppendorf.

Microbiological safety cabinet:

Add 1ml of 70% EtOH to seeds and shake for 2 minutes.

Remove and add 1ml of sterilising solution, shake for 6 – 8 minutes.

Remove and add 1 ml of 70% EtOH and shake for 2 minutes.

Wash 5 times with sterile dH_2_O.

Top up the Eppendorf with sterile dH_2_O, wrap in parafilm and tin foil and store in the fridge for 2 days.

Pipette 12 seeds in 3 rows of 4 seeds onto prepared ½ MS plates.

Place vertically in growth cabinet set at 20°C for a 12-hour light/dark cycle.

**½ MS Agar 600ml:**

- 1.32g Murashige and Skoog powder
- 600 ml dH_2_O
- 4.8g Agar

Add the MS powder to the water and adjust the pH to 5.7.

Add agar and adjust the pH to 6.72.

Autoclave at 121°C for 15 minutes.

Microbiological safety cabinet:

Pour 50ml into square petri dishes and leave to set.

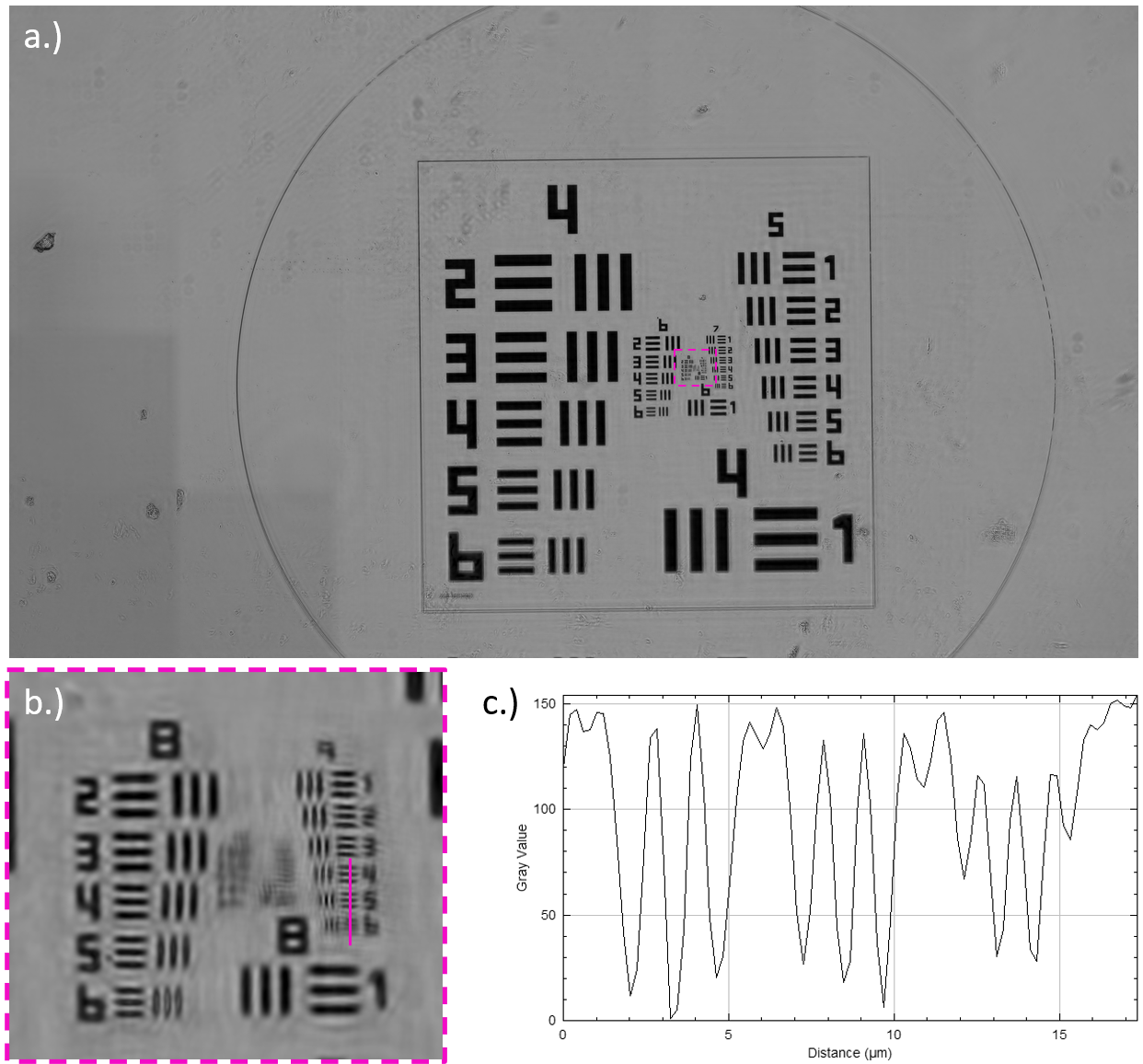
USAF target imaging and resolution measurements:

Figure S1: US Air Force test target imaged with the red (636 nm) LEDs. a.) shows the full FOV of the test target. b.) is a digital zoom ROI of the centre of a.) with a line through the smallest line pairs of group 9. c.) is the resulting line plot profile of the three smallest line pairs using FIJI. [20]

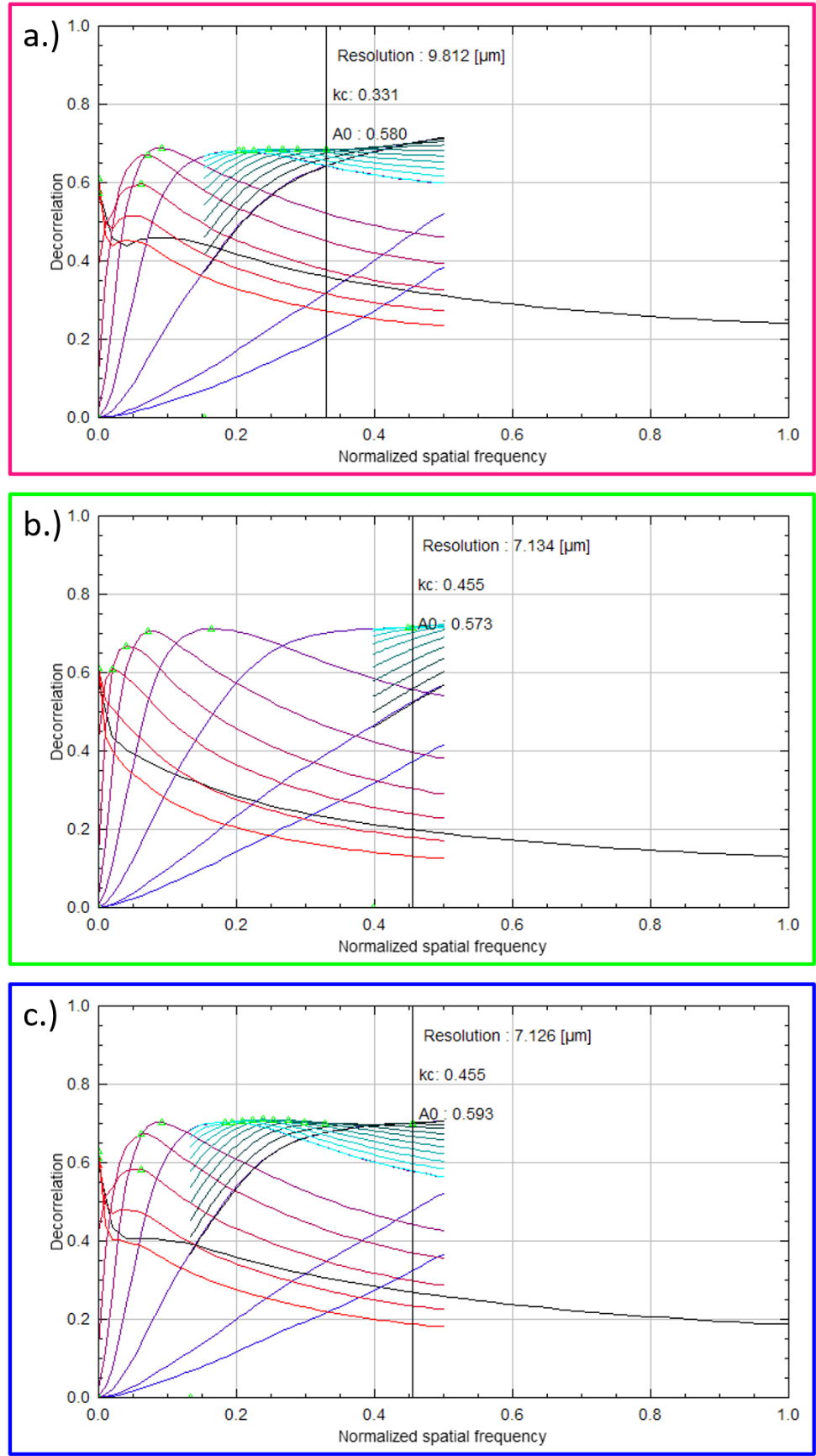
Image decorrelation Analysis:

Figure S2: Decorrelation analysis plots of incoherent images imaged with a.) red (636 nm) LEDs, b.) green (523 nm) LEDs, and c.) blue (470 nm) LEDs.

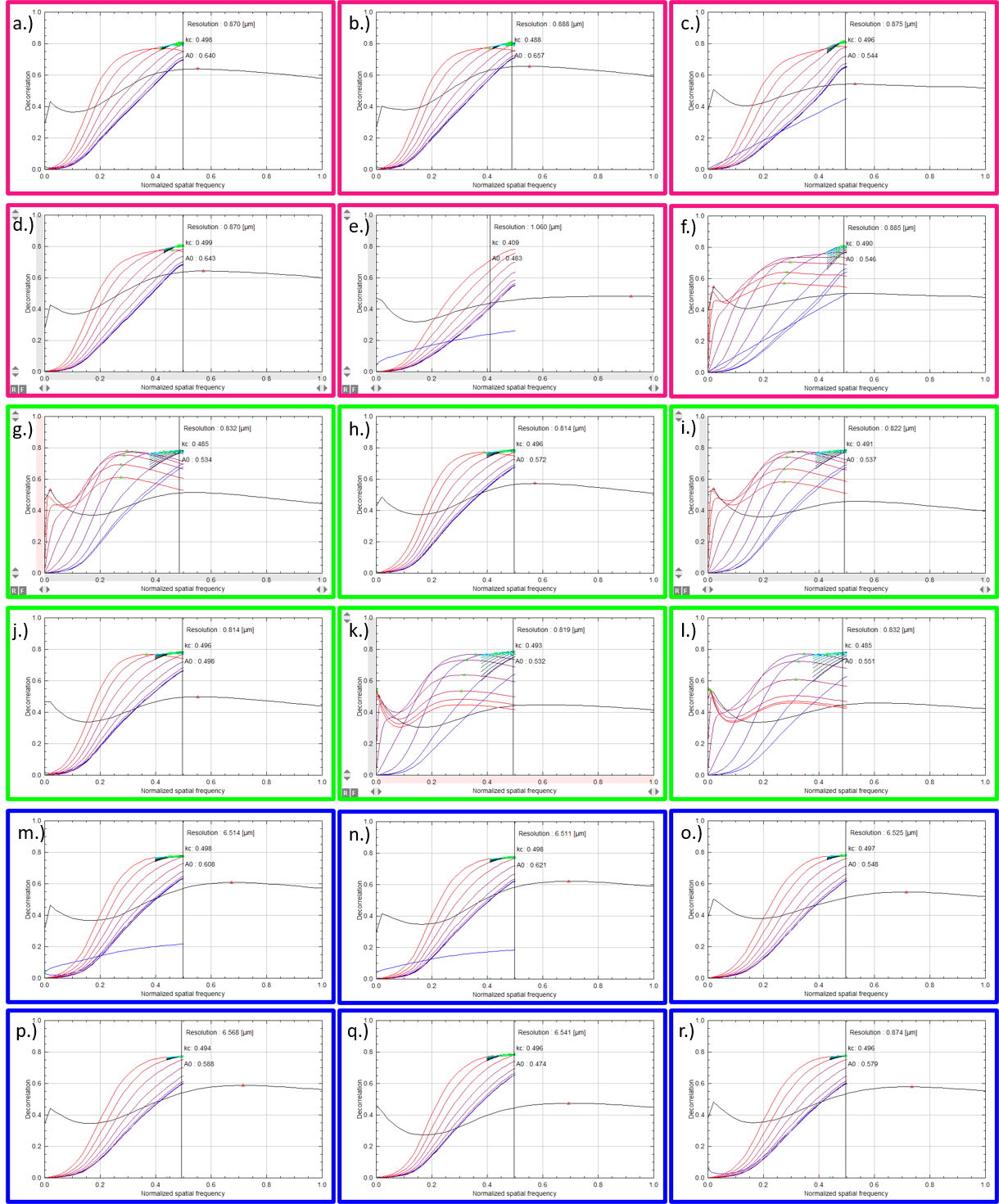
Figure S3: Image decorrelation analysis of FPM amplitude images illuminated with red (636 nm), green (523 nm) and blue (470 nm) LEDs. a.) – f.) are the images using red LEDs, g.) – l.) are the images using green LEDs and m.) – r.) are the images using blue LEDs.

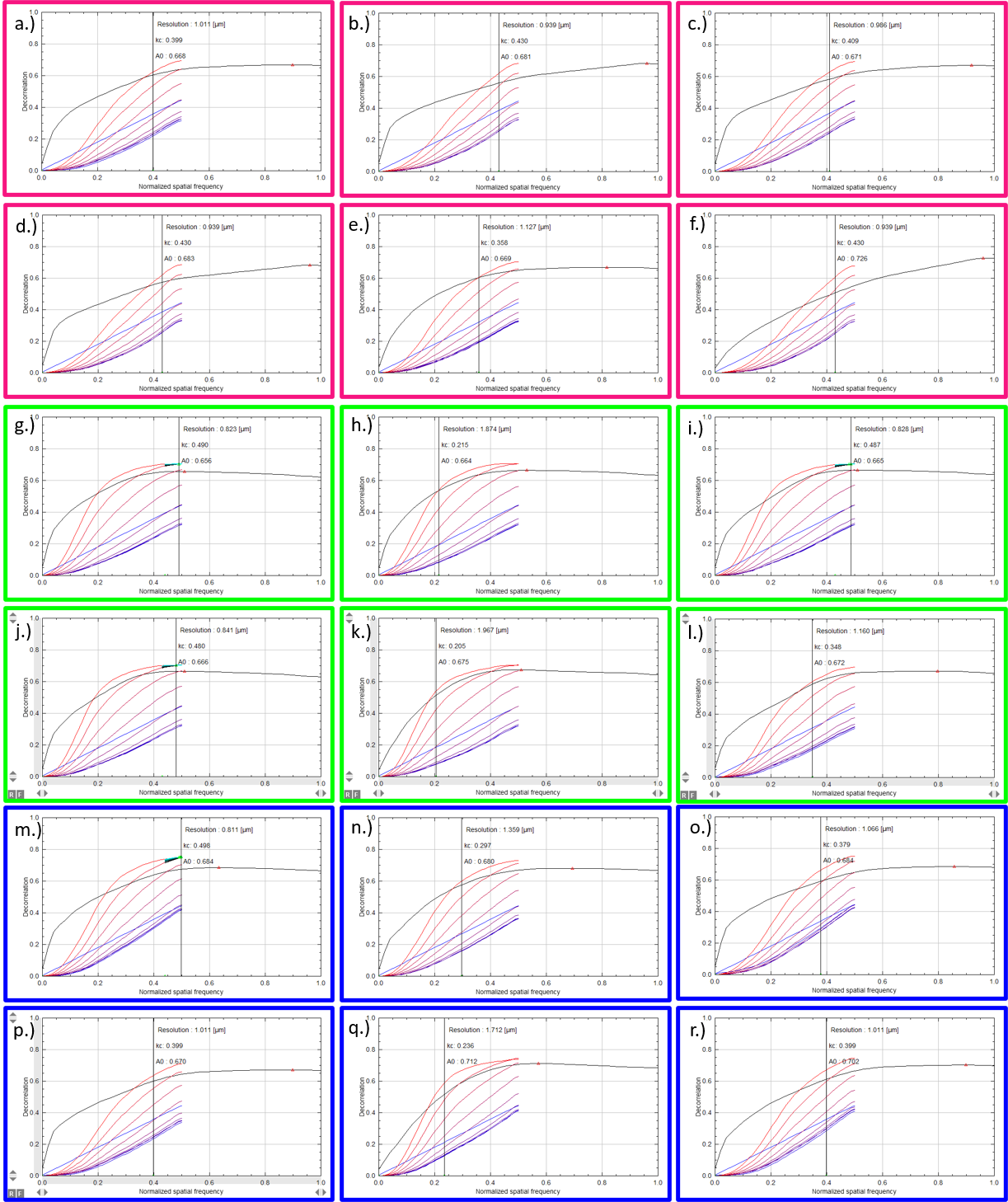
Figure S4: Image decorrelation analysis of FPM phase images illuminated with red (636 nm), green (523 nm) and blue (470 nm) LEDs. a.) – f.) are the images using red LEDs, g.) – l.) are the images using green LEDs and m.) – r.) are the images using blue LEDs.

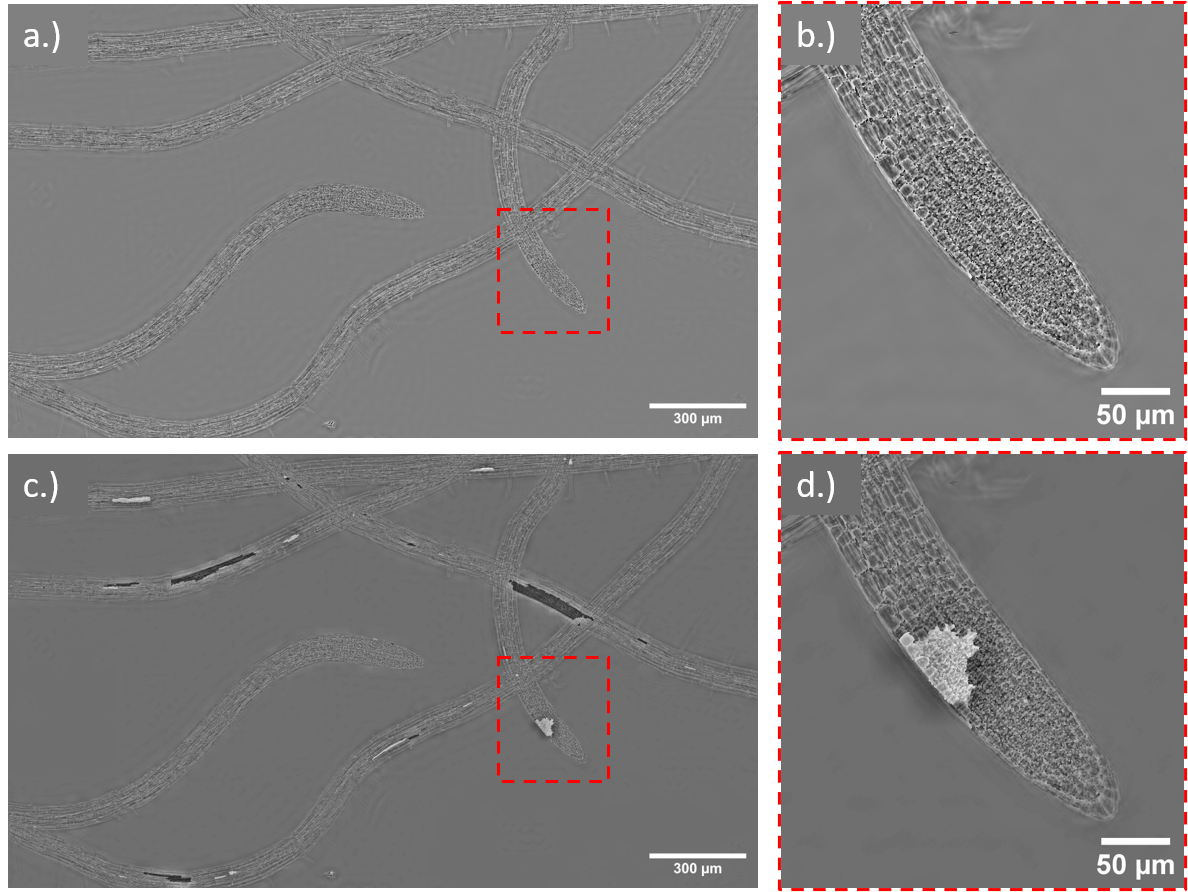

Figure S5: Phase unwrapping artefacts in *Arabidopsis thaliana* image. a.) is the FPM phase image with wrapping artefacts. c.) is the phase image after going through the phase unwrapping algorithm. [27] b.) and d.) are digital zoom ROIs of the root tip to highlight that the phase wrapping appears uniform across b.) however the artefact is contained to just one patch in d.).
